# A hierarchy of processing complexity and timescales for natural sounds in human auditory cortex

**DOI:** 10.1101/2024.05.24.595822

**Authors:** Kyle M. Rupp, Jasmine L. Hect, Emily E. Harford, Lori L. Holt, Avniel Singh Ghuman, Taylor J. Abel

## Abstract

Efficient behavior is supported by humans’ ability to rapidly recognize acoustically distinct sounds as members of a common category. Within auditory cortex, there are critical unanswered questions regarding the organization and dynamics of sound categorization. Here, we performed intracerebral recordings in the context of epilepsy surgery as 20 patient-participants listened to natural sounds. We built encoding models to predict neural responses using features of these sounds extracted from different layers within a sound-categorization deep neural network (DNN). This approach yielded highly accurate models of neural responses throughout auditory cortex. The complexity of a cortical site’s representation (measured by the depth of the DNN layer that produced the best model) was closely related to its anatomical location, with shallow, middle, and deep layers of the DNN associated with core (primary auditory cortex), lateral belt, and parabelt regions, respectively. Smoothly varying gradients of representational complexity also existed within these regions, with complexity increasing along a posteromedial-to-anterolateral direction in core and lateral belt, and along posterior-to-anterior and dorsal-to-ventral dimensions in parabelt. When we estimated the time window over which each recording site integrates information, we found shorter integration windows in core relative to lateral belt and parabelt. Lastly, we found a relationship between the length of the integration window and the complexity of information processing within core (but not lateral belt or parabelt). These findings suggest hierarchies of timescales and processing complexity, and their interrelationship, represent a functional organizational principle of the auditory stream that underlies our perception of complex, abstract auditory information.

## Introduction

Humans encounter a wide and diverse range of sounds in daily life, requiring rapid categorization to support efficient behavior. Is a cell phone vibrating in the other room or did a bee get inside? Was that a whisper or just the wind? The brain recognizes sounds with vastly different acoustic signatures as functionally equivalent (e.g., a laugh and scream are human vocalizations) and also differentiates acoustically similar sounds across incongruent classes (e.g., a man humming and a flute playing the same note). The cortical mechanisms responsible for our ability to differentiate and identify sounds according to their spectrotemporal acoustic properties has been studied extensively as auditory categorization (1–4). There is widespread consensus for a hierarchical organization of auditory cortex, with primary areas representing acoustic features and downstream regions encoding progressively more abstract representations better aligned with category-level representations. Yet, the degree of complexity of auditory cortical representations has been difficult to quantify objectively, limiting our ability to describe and account for the progression of information throughout this hierarchy.

Prior work suggests early auditory responses can be explained by their responsivity to sounds’ spectrotemporal features and modulations thereof (5–8). A spectrotemporal modulation (STM) tuning framework has been applied to both primary and downstream auditory cortex to characterize encoding of these features in the context of speech (9, 10). Furthermore, STM models have been used to reconstruct speech (11) and natural sounds from neural responses (12). However, the majority of acoustic sensory input we receive falls within only a subregion of STM space (13, 14), which may partially explain why this approach has been unable to produce a fully generalizable model of neural responses outside of primary auditory cortex (2, 5, 15–20). More importantly, STM features measure acoustic properties and are thus not well-suited to describe responses in higher-order auditory cortex, which responds to sound categories such as vocalizations (21) and music (16), exemplars of which may have highly distinct spectrotemporal features.

Recently, task-optimized deep neural network (DNN) models have been used to explore representations throughout auditory cortex, generating compelling evidence that supports a non-linear hierarchically-organized auditory network (15, 17, 18, 22–25) that transforms incoming acoustic input (e.g. speech, music, environmental sounds) into salient, behaviorally-relevant representations. Critically, commonly-used feedforward DNN architectures that incorporate non-linear transformations at each layer result in stimulus representations that become increasingly complex with DNN layer depth (26). In the case of a DNN trained to classify sounds into semantic categories, this transformation approximates a continuum from acoustic feature encoding in shallow layers to abstract, semantic representations in deep layers (15, 27, 28). This framework can be used to quantitatively estimate representational complexity, i.e., the complexity of feature representations, across auditory cortex.

Here, in the context of epilepsy surgery, we acquired intracerebral electrophysiology from 20 patient-participants as they listened to a rich set of natural sounds that spanned multiple categories, including speech and non-speech vocalizations, music, animal vocalizations, and environmental sounds (16). We extracted sound features from different layers within a sound-categorization DNN and built encoding models to predict neural responses throughout auditory cortex. This task-optimized DNN was trained to evaluate a sound’s spectrogram and classify it within a broad hierarchical taxonomy (e.g., human vocalizations such as speech and laughter, music genres and instruments, types of mechanical sounds, and so on) and thus was well-suited to explore natural sound encoding in human auditory cortex. Across recording sites, we used these encoding models to assess the relationship between neural representations and the layers of the DNN. The representational complexity was inferred by the layers of the DNN that best explained the neural responses, with deeper layers corresponding to more complex representations; we then characterized how complexity varied across and within core, belt, and parabelt regions. Lastly, we used these encoding models to estimate the temporal windows over which each recording site integrates information and then related the integration windows to both representational complexity and anatomical position within the auditory cortical hierarchy. By characterizing how complexity and timescales vary (and covary) across auditory cortex, we elucidate functional organizational principles underlying the processing of natural sounds in auditory cortex.

### Methods

#### Patient-Participants

Data were collected from 20 adult and pediatric patients (age 10 – 25 years old, 7 females) with epilepsy undergoing in-patient stereo-electroencephalography (sEEG) monitoring at UPMC Children’s Hospital of Pittsburgh.

Each patient had between 6-20 electrode trajectories, with each trajectory containing between 8-18 electrode contacts, which will henceforth be referred to as channels. sEEG electrodes were implanted with a robot-assisted frameless stereotactic technique, as previously described (43). All electrode locations were selected based purely on clinical considerations. The research protocol was approved by the University of Pittsburgh Institutional Review Board (STUDY20030060). Further demographic information can be found in Table 1.

**Table 1.** Participant demographic and clinical information.

| ID | Age (yrs) | Sex | Handedness | Language dominant hemisphere | Epilepsy onset (years) | Seizure locus | Epilepsy etiology | sEEG laterality | Electrode # (auditory responsive/all) |
| --- | --- | --- | --- | --- | --- | --- | --- | --- | --- |
| P1 | 19 | Female | Right | Unknown | 14 | Right SMA | Lesional | Bilateral | 55/106 |
| P2 | 13 | Female | Right | Unknown | 3 | Bilateral MTL | Unknown | Bilateral | 27/104 |
| P3 | 14 | Male | Right | Unknown | 13 | Tumor | Tumoral | Left | 47/54 |
| P4 | 18 | Male | Right | Left | 13 | Left multifocal | Autoimmune | Bilateral | 39/127 |
| P5 | 14 | Female | Left | Unknown | 14 | Right MTL | Unknown | Right | 24/125 |
| P6 | 15 | Male | Right | Unknown | 14 | Right posterior MTL | FCD | Bilateral | 55/117 |
| P7 | 15 | Female | Right | Left | 13 | Left MTL | MTS | Left | 63/119 |
| P8 | 10 | Male | Right | Unknown | 8 | Bilateral multifocal | Autoimmune | Bilateral | 24/120 |
| P9 | 16 | Male | Right | Unknown | 0 | Right frontal | Unknown | Right | 51/226 |
| P10 | 19 | Male | Ambidextrous | Left | 17 | Left MTG/STG | Tumoral | Left | 23/56 |
| P11 | 16 | Male | Left | Right | 1 | Right parietal + cingulate | FCD | Bilateral | 71/256 |
| P12 | 17 | Male | Right | Unknown | 14 | Right posterior MTL | FCD | Right | 34/164 |
| P13 | 19 | Male | Right | Unknown | 2 | Right STG | Unknown | Right | 37/205 |
| P14 | 16 | Male | Left | Unknown | 12 | Left frontal | Unknown | Bilateral | 25/256 |
| P15 | 22 | Male | Right | Right | 12 | Right multifocal | Unknown | Bilateral | 51/243 |
| P16 | 17 | Female | Right | Left | 15 | Left multifocal | Unknown | Bilateral | 44/200 |
| P17 | 19 | Female | Right | Unknown | 12 | Left MTL | MTS | Left | 41/151 |
| P18 | 16 | Male | Ambidextrous | Unknown | 12 | Left frontal | Unknown | Bilateral | 8/218 |
| P19 | 25 | Male | Left | Left | 2 | Left parietal | Lesional | Left | 72/126 |
| P20 | 13 | Female | Right | Unknown | 11 | Left MTL | Tumoral | Left | 20/172 |

#### Data collection

Patients performed an auditory one-back task while listening to a well-characterized stimulus set of natural sounds (16) consisting of 165 two-second clips of speech (English and non-English), human and animal vocalizations, music, and other environmental sounds (see Fig. 1A). The 165 stimuli were presented in random order, with a random 20% chosen to be followed by an immediate repeat (i.e., a one-back target), and an interstimulus interval that varied randomly from a uniform distribution ranging from 1-2 seconds. This resulted in 198 trials, which were split into two blocks. The process that generated this stimulus order was repeated three times, resulting in six blocks and 594 trials. Patients indicated the presence of a one-back target using a button box (RT Box, model v6). The experiment was run using Psychtoolbox-3 and custom MATLAB code.

**Fig. 1.**
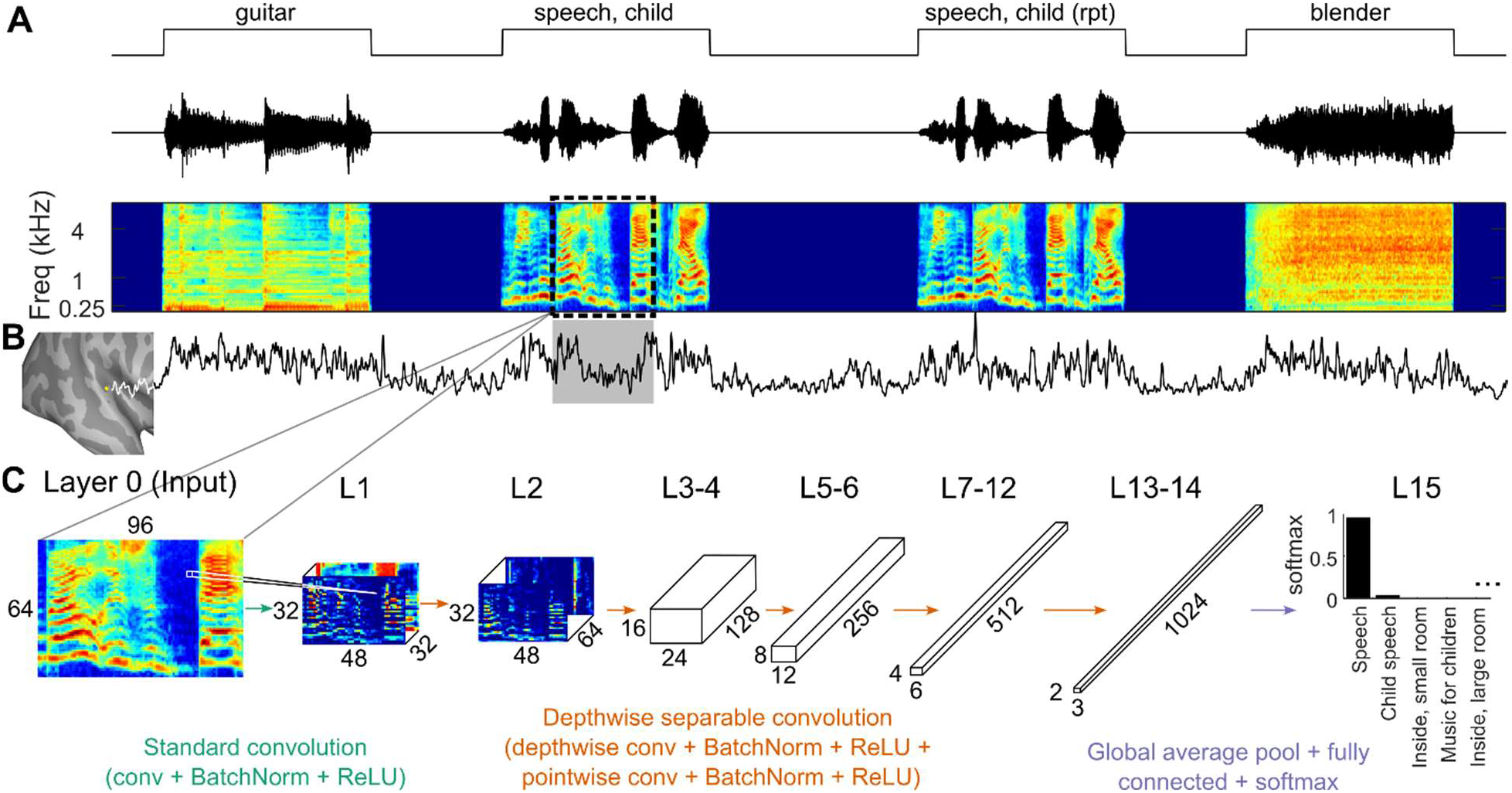
Methods. (A) Patients performed an auditory 1-back task using Natural Sounds stimuli. The dashed black box in the auditory spectrogram represents the 975 ms input window for the DNN (see panel C). (B) Broadband high gamma activity (HGA) from an example channel. (C) YAMNet deep neural network model architecture. Arrow colors represent different blocks of DNN layer operations. Depthwise separable convolutions were also used between the grouped layers in the figure (L3-4, L5-6, L7-12, and L13-14). Using this pre-trained DNN, layer activations for each stimulus were extracted and used to build encoding models to predict HGA.

Neural data were sampled at 1000 Hz using a Ripple Grapevine Nomad neural interface processor (model R02000), with line noise notch filters at 60, 120, and 180 Hz. Audio was split using a distribution amplifier (Rolls model DA134), with one stream presented to the patient at a volumed deemed loud but comfortable via Etymotic ER-3C insert earphones, and a second stream recorded synchronously with neural data using a Ripple Digital/Analog IO box (model R02010-0017) and sampled at 30 kHz. Digital stimulus triggers marking the onsets and offsets of each stimulus were sent using a Measurement Computing Data Acquisition device (model USB-1208FS) and were recorded via the IO box.

#### Preprocessing

Data were common average reference filtered and epoched to include 1000 ms pre-stimulus onset to 3000 ms post-stimulus onset. Stimulus onsets were determined via cross-correlation between the original stimulus files and the audio recorded synchronously with neural data. Data in each channel was then z-scored relative to the baseline signal calculated across all trials.

High gamma activity (HGA, 70-150 Hz) was then extracted in the following way. For each channel, data was forward- and reverse-filtered (Butterworth, sixth order) in eight different bands, with center frequencies and bandwidths logarithmically spaced between 70-150 Hz and 16-64 Hz respectively. The analytic signal amplitude for each band was extracted using the Hilbert transform, and the resultant signal was z-scored across all trials relative to a baseline period of -900 to -100 ms relative to stimulus onset.

The first and last 100 ms of the 1000 ms baseline period were excluded to avoid contamination from edge effects and any rapid onset neural responses, respectively. After z-scoring, the signal was averaged across frequency bands and then down-sampled to 100 Hz. Lastly, the signal was z-scored across trials once more, again using a baseline window of -900 to -100 ms. This measure represents HGA, which was averaged across all presentations of a given stimulus for each channel (termed a channel’s stimulus response).

#### Anatomy

The pipeline for determining channel locations and generating anatomy plots consisted of the following steps. For each patient, a cortical surface was reconstructed from a pre-operative T1-weighted MRI using Freesurfer (29). The output files were then imported into Brainstorm (30), a third-party tool developed in MATLAB that was used for all subsequent steps in this anatomy pipeline. A post-operative CT was co-registered to the MRI and then used to mark individual channel locations. Non-linear MNI normalization was then performed in Brainstorm, which internally uses SPM12 for the procedure (31).

Lastly, the Julich Brain (v3.0) volumetric atlas (32) was imported for region-of-interest (ROI) analysis, using the MNI reverse field deformation to transform the atlas into the patient-specific spaces. The core/belt/parabelt taxonomy was used with the following definitions: core – Te1.0 and Te1.1; medial belt – TeI; lateral belt – Te1.2, Te2.1, and Te2.2; parabelt – Te3, STS1, and TPJ (33–36). The combination of these regions constituted our definition of auditory cortex. Due to sparse coverage in medial belt, this region was excluded from most analyses.

To define a channel’s precise location within an ROI along anatomically relevant dimensions (e.g., in the analysis for Fig. 3), we calculated a set of three orthogonal axes for each of core, lateral belt, and parabelt (one for each hemisphere, resulting in six total sets of axes). The axes for parabelt were determined using eigendecomposition. Using a three-dimensional point cloud of MNI voxels that belong to parabelt in the Julich atlas, we calculated the eigenvectors of this (demeaned) point cloud’s covariance matrix. The eigenvector associated with the largest eigenvalue points along the axis of highest variance, which in this case is oriented from posterior to anterior along the length of superior temporal gyrus. The other two eigenvectors point roughly from ventral to dorsal and from medial to lateral. Each axis was then defined as a line pointing along a given direction and centered at the point cloud’s centroid. The edges of an axis were defined as 0-1 and were found by projecting the ROI’s voxels to the axis and finding the furthest points from the centroid. Eigendecomposition was also used to find the primary axis in core; the eigenvector with the largest eigenvalue points from posteromedial to anterolateral along Heschl’s gyrus. The secondary axis was calculated to run across Heschl’s gyrus by finding a vector perpendicular to the primary axis and parallel to the supratemporal plane (STP, defined as Te1.0, Te1.1, Te1.2, Te2.1, Te2.2, TeI, and TI); the tertiary axis was thus perpendicular to STP. In lateral belt, the core axes directions were used, with the origin shifted to the lateral belt centroid. Each channel in a given ROI was then projected to that ROI’s axes and normalized using the aforementioned 0-1 scaling.

MNI normalization for one patient produced abnormal results due to a prior stroke in frontal lobe, mapping channels to incorrect locations on the MNI brain and producing incorrect ROI labels (since ROI labeling is dependent on the MNI transformation). For this patient, ROI labels were manually corrected to include in ROI analysis; his channels were not displayed on any MNI brain surface plots.

#### Encoding models

Encoding models were used to predict neural responses to novel auditory stimuli. First, HGA was averaged across all presentations of a stimulus. We built models to predict two different measures of HGA: 1) the mean HGA stimulus response across a single 975 ms window per stimulus, starting at 480 ms after stimulus onset, which we call the *long-window model*, and 2) the short-window HGA response, in which HGA stimulus responses were averaged over shorter windows to investigate temporal encoding characteristics (see *Short-window encoding models*). For the long-window model, the window size (975 ms) was chosen to match the input window for the deep neural network under consideration here, and we chose to analyze responses starting at 480 ms to avoid any generic acoustic onset effects and focus solely on sustained responses (37–39). The same temporal window (480-1455 ms) was used to calculate the layer activations and the mean HGA.

Using the MATLAB package *glmnet* (40, 41), we built L1-regularized regression models for each channel. (See the following section, *Stimulus features*, for a detailed description of encoding model inputs.) A nested cross-validation was used, with the inner cross-validation (10-fold) used to select the regularization parameter lambda; the outer cross-validation (5-fold) was used to test the resultant model by generating HGA predictions on stimuli that were held out of the training set. Model accuracy was assessed by calculating the coefficient of determination (R2) between observed and predicted HGA. This measure describes the fraction of the HGA variance explained by a model and is calculated by summing the squared residuals, dividing this by the sum of squared residuals for a baseline model that only uses an intercept term (i.e., all responses are modeled as the mean HGA response), and subtracting this ratio from one. When characterizing which DNN layers produced the best models for each channel, we calculated the weighted mean of the R2 curve across layers, which we refer to as the weighted DNN layer. Since R2 can sometimes produce negative values (if a model performs worse than the baseline intercept model), and a weighted mean requires all weights to be nonnegative, we set any negative R2 values to zero when calculating the weighted DNN layer.

#### Stimulus features

Stimulus features were generated from the DNN model YAMNet (27), inspired by the success of image classification models such as AlexNet (42). This machine learning model was trained to predict sound classes on thousands of hours of labeled audio taken from AudioSet (28), a large-scale library of sounds scraped from YouTube that have been manually tagged with sound categories of a hierarchical ontology of classes. For example, a given sound might be simultaneously tagged as Music, Soul music, Singing, and Female singing. Note that this DNN model was built and trained by an unaffiliated machine learning research team, was held static in this research (i.e., no further training was performed), and did not have access to any neural data, including the data used in this study.

The YAMNet model consists of 16 layers (depicted in Fig. 1C), starting with an input layer 0, which accepts mel-spectrograms of sounds, and layer 1 that performs standard 2D convolutions. Layers 2-14 are depthwise separable convolutional layers, where each layer consist of two convolutions: first, a grouped depthwise convolution is performed, where each filter is associated with and learned on a single channel (here, channel refers to the third dimension, or depth, in each DNN layer). The second convolution in a depthwise separable layer is a pointwise convolution, where each filter is 1 x 1 x number of channels, allowing the layer to mix information across channels. Each individual convolution is followed by a batch normalization and a rectified linear unit (ReLU). The final layer consists of a global average pooling layer, a fully connected layer, and a softmax output that generates a probability distribution over the discrete sound classes.

Each of the 165 stimuli was provided as input to the YAMNet model, and activations across all nodes within a given layer were calculated. These layer activations served as stimulus features; repeating this process for all 16 layers thus produced 16 unique feature sets, numbered 0 (the input spectrogram) to 15 (the probability that the sound belongs to each semantic class).

The YAMNet model has not been trained to mimic human neural representations of sound features. Rather, the training objective function dictates that the model iteratively adjusts its weights to solve the task while minimizing the cross-entropy, a measure of the distance between the output class probabilities and the ground truth labels. Any similarities that arise between internal model representations and human neural responses is thus an emergent property of the model. The DNN learns to extract an optimal set of stimulus features for audio classification and can thus be viewed as a tool for data-driven feature extraction.

Furthermore, the model imposes a natural and interpretable gradient from early layers that represent low-level acoustic properties, through middle layers with more complex acoustic representations, to the deepest layers that represent wholly abstract stimulus features (i.e., the sound category). Each layer performs a nonlinear transformation, introducing further complexity in the stimulus representation with increasing layer depth. We can leverage these evolving stimulus representations to quantify the degree of representational complexity present at different cortical sites by identifying the DNN layers that are best able to explain the neural responses. Theoretically, an encoding model that can predict neural responses in human auditory cortex with high fidelity using the activations (i.e., the responses) within a YAMNet hidden layer would demonstrate that there is shared information between the two systems.

#### Short-window encoding models

Short-window models were used to characterize temporal encoding properties and were built only for channels that achieved a peak R2 > 0.1 (i.e., maximum R2 across DNN layers) in the long-window analysis. First, we determined the shortest window over which to average HGA that still contained meaningful stimulus information in the context of DNN stimulus features, which we refer to as the encoding window (43). Specifically, this was determined as follows for a given channel. The DNN layer that produced the peak R2 in the long-window analysis was selected, and an encoding model was built to predict HGA averaged over a 20 ms window centered on the leading edge of the stimulus feature window (i.e., if we define the stimulus feature window as spanning from -975 to 0 ms, HGA was averaged from -10 to 10 ms). If this encoding window was too short to produce a sufficiently good model (R2 > 0.1), then the process was iterated with a wider window until the R2 > 0.1 threshold was satisfied. A total of 11 candidate encoding windows were explored, essentially logarithmically spaced (with rounding to the nearest 10 ms, since HGA was down-sampled to 100 Hz) up to 960 ms. These candidate windows were 20, 40, 60, 80, 120, 160, 240, 340, 480, 680, and 960 ms. If none of these encoding windows produced a model with R2 > 0.1, then that channel was discarded for all short-window analysis. Once an encoding window was selected, models were built and evaluated using this same window for all 16 DNN layers.

For the short-window analysis, eight observations were extracted for each of the 165 stimulus responses; the earliest of these was positioned with the leading edge of the stimulus feature window at 960 ms relative to stimulus onset, with the other seven observations spaced 80 ms apart up to 1520 ms, resulting in a total of 1320 observations.

#### Statistical analyses

Most statistical tests, including ROI analyses, were performed using linear mixed effects (LME) modeling. All LME models included hemisphere and ROI as fixed effects and patient as a random effect (intercept only).

## Results

Across 20 patients, there were a total of 3145 intracerebral recording sites, of which 811 were auditory-responsive, defined as a statistically significant difference between HGA stimulus responses compared to baseline (two-sample t-test, p < .01, false discovery rate corrected). Of those, 755 channels exhibited increased responses, while 56 channels showed decreased responses. There were 386 channels localized to auditory cortex, with 303 of these exhibiting auditory responses (core: 91 auditory-responsive/93 total channels, medial belt: 9/9, lateral belt: 105/106, and parabelt: 98/178). All auditory-responsive channels in auditory cortex exhibited increased responses. Due to the sparse coverage within medial belt, this region was excluded from all ROI analyses.

First, we sought to establish whether a DNN trained to categorize sounds possessed similar representations to channels throughout human auditory cortex. To this end, we built long-window (975 ms) encoding models to predict broadband HGA responses for all auditory-responsive channels using layer activations from the deep neural network YAMNet. This pipeline is shown for three example channels in Fig. 2A-D. Note that the analysis window (gray box, Fig. 2B) is delayed from stimulus onset

**Fig. 2.**
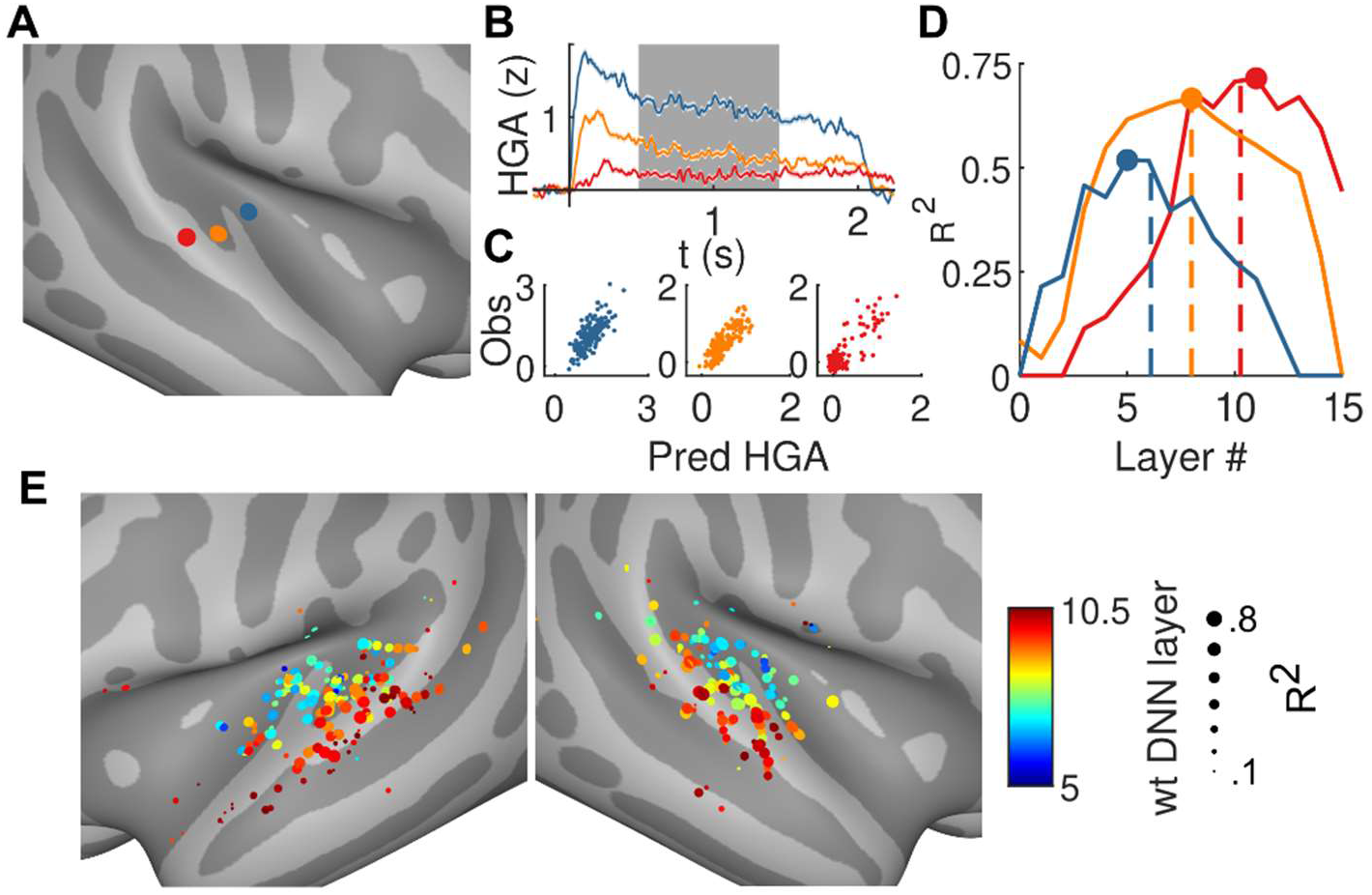
Long-window encoding model results. (A) Example channels in core (blue), lateral belt (orange), and parabelt (red) used for panels B-D. (B) Mean HGA (±1 SEM). Gray box shows analysis window for long-window models (see Methods). (C) Predicted vs. observed responses for the best-performing encoding models. (D) Model accuracies across DNN layers. Points show best (peak) model plotted in (C), and dashed lines show the weighted DNN layer, which is the weighted mean of each curve. (E) Encoding model results across patients and channels. Neural prediction accuracy for the best model is shown by marker size. Color represents the weighted DNN layer (the dashed lines from panel D).

by 480 ms to avoid any onset responses, so that analyses could focus solely on sustained responses. Fig. 2D shows encoding accuracy across all layers of YAMNet for the three example channels. Note that the blue, orange, and red channels (which correspond to core, belt, and parabelt) are best predicted by shallower, middle, and deeper layers respectively. Fig. 2C shows predicted versus observed responses for the peak models of each channel (i.e., the dots in Fig. 2D).

Neural responses throughout auditory cortex could be predicted with high accuracy, achieving R2 values up to 0.81, while most auditory-responsive channels outside of auditory cortex could not be predicted well using these encoding models. Of the 303 auditory-responsive channels in auditory cortex, 255 channels (84%) achieved a peak R2 (i.e., maximum R2 across DNN layers) greater than 0.1. In contrast, only 14% (69/508) of auditory-responsive channels outside of auditory cortex achieved a peak R2 over 0.1, suggesting that most responses outside of auditory cortex do not encode the stimulus features captured by the DNN. Lastly, channels with decreased stimulus responses relative to baseline (all of which were outside of auditory cortex) could not be predicted well with these encoding models; all 56 such channels achieved a peak R2 less than 0.01. Within auditory cortex, peak R2 values were lower in parabelt compared to core (p = 3.8x10-3) and lateral belt (p = 4.3x10-4) regions, even when controlling for auditory responsiveness (LME model with R2 as the response variable). This effect is explained by the fact that only 67% of auditory-responsive parabelt channels achieved a peak R2 > 0.1, compared to 93% of core channels and 94% of lateral belt channels.

Rerunning the LME model using only channels with peak R2 > 0.1 (250 total channels in core, lateral belt, and parabelt) yielded no significant differences between these three ROIs. In other words, core and lateral belt regions appear to broadly encode DNN features, while only a subset of parabelt does, potentially due to the functional heterogeneity within this region. However, of the recording sites that do encode these features, there are no systematic differences in encoding performance between these three regions.

We then estimated the representational complexity, which we define as the complexity of features encoded by each channel. This was quantified as the DNN layer that produced the most accurate predictions of neural responses, with deeper layers having undergone more nonlinear transformations and thus corresponding to more complex features. While this could be done by selecting the DNN layer that maximizes R2 (24), this approach is susceptible to noise. Instead, we calculated the weighted DNN layer, defined as the weighted mean of R2 across DNN layers (i.e., the center of mass of the curves in Fig. 2D, shown as dashed lines). Fig. 2E shows weighted DNN layers and peak R2 values plotted across patients and channels, revealing a relationship of increasing representational complexity in higher-order compared to lower-order auditory cortex. This effect was statistically validated using an LME model and channels with peak R2 > 0.1. Controlling for auditory responsiveness, core mapped to shallower layers relative to lateral belt (p < 10-5) and parabelt to deeper layers (p = 0.015; also see Fig. 5, top panel).

There was also a hemispheric effect, with left hemisphere mapping to deeper DNN layers than right hemisphere when controlling for ROI (p = 0.012).

In addition to complexity differences between ROIs, we hypothesized that complexity gradients existed within each ROI along specific axes (see Methods for descriptions on how each axis was calculated). In core, the primary axis under consideration was oriented from posteromedial to anterolateral (PM-AL, Fig. 3A). An orthogonal secondary axis that was parallel to the STP, as well as a tertiary axis perpendicular to STP, were also investigated (not shown). In lateral belt, axes with the same orientation as core were used, with the origin shifted to the lateral belt centroid. In parabelt, the three axes pointed from posterior to anterior (P-A) along the length of superior temporal gyrus (STG), from ventral to dorsal (V-D), and from medial to lateral (Fig. 3B, M-L axis not shown). Channels belonging to a given ROI were projected to each of that ROI’s axes, and correlations were calculated between these positions and representational complexity (i.e., weighted DNN layer). In both core and lateral belt, representational complexity increased as a function of position from posteromedial to anterolateral, but only in the right hemisphere (Fig. 3A). No significant relationship was observed along the secondary or tertiary axes (parallel to and perpendicular to STP, respectively). In parabelt, complexity increased moving anteriorly as well ventrally (Fig. 3B). Again, this relationship was observed only in the right hemisphere. Position along the M-L axis was not correlated with complexity in parabelt.

**Fig. 3.**
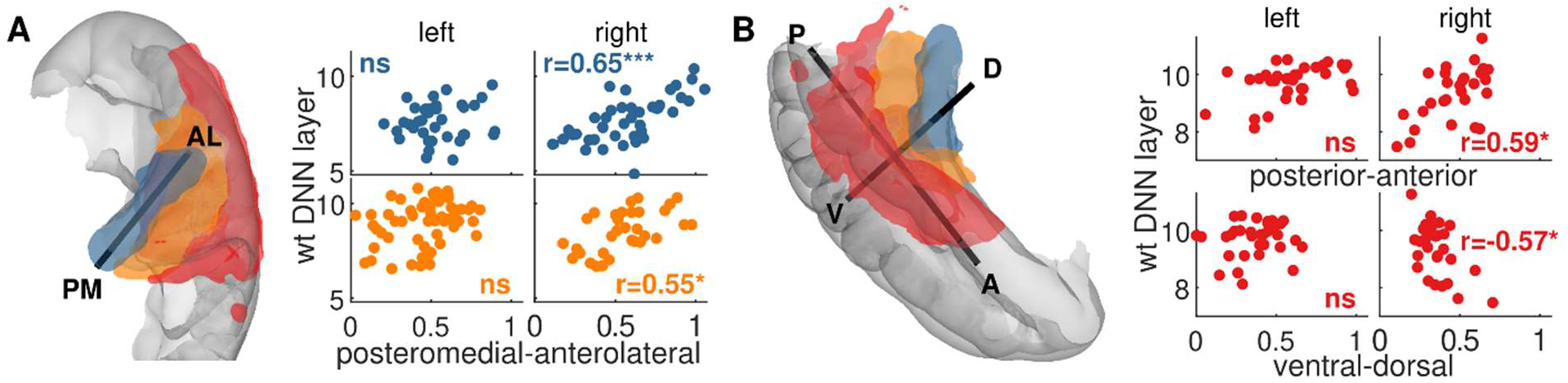
Complexity gradients within ROIs. (A) A gradient of increasing representational complexity (indexed by weighted DNN layer) was found along a posteromedial-anterolateral axis in both core and lateral belt, but only in the right hemisphere. This axis was defined using the best fit line through core voxels. For lateral belt, we used an axis with the same direction but shifted to the lateral belt centroid. (B) Representational complexity gradients were also found in parabelt along the posterior-anterior and ventral-dorsal axes, with complexity increasing in the anterior and ventral directions. Again, this relationship was only observed in the right hemisphere. Results are across all patients and channels with long-window R^2^ > 0.1. *** p < .001, ** p < .01, * p < .05, Bonferroni corrected.

We then tested whether these within-ROI complexity gradients were best explained by a linear or sigmoidal relationship. A linear relationship would demonstrate a smoothly varying degree of complexity along the axis, while a sigmoidal relationship might indicate a more step-like transition, which would be consistent with homogenous-complexity subregions within an ROI. This test was performed by fitting both linear and sigmoid models to the channel position versus weighted DNN layer relationship (i.e., the scatter plots in Fig. 3), and then comparing the two models using the Akaike information criterion (AIC).

The sigmoid model contained four parameters that described the lower and upper asymptotes, the x-position of the midpoint between asymptotes, and the slope at that midpoint (i.e., how step-like the transition is). We required the x-position to fall between 0 and 1 but placed no other constraints when fitting the sigmoid model. We found that a linear model better explained these relationships than a sigmoidal one in right core (PM-AL, relative likelihood of sigmoidal compared to linear = 0.135) as well as parabelt (P-A relative likelihood = 2.41x10-4; V-D relative likelihood = 0.139). In contrast, the complexity gradient within right lateral belt was better explained by a sigmoidal relationship (PM-AL, relative likelihood of linear compared to sigmoidal = 0.217). In summary, representational complexity appeared to vary more linearly in right core and parabelt but was more step-like in right lateral belt.

Next, we sought to characterize temporal encoding properties throughout auditory cortex for all channels whose long-window models achieved peak R2 > 0.1. To quantify these dynamics, we examined the *integration window*, which refers to the temporal extent over which a neural population represents stimulus information (43). According to this definition, changes to the stimulus outside this window do not produce changes in the neural response, while changes inside of it do. To estimate the integration window for a given channel, we first characterized the *encoding window*, which refers to the window over which neural response modulations contain meaningful information (43). While integration windows and encoding windows describe different properties, they are intrinsically linked; see Theunissen and Miller (43) for more detail.

To carry out this analysis, we built short-window models, in which broadband HGA was averaged over shorter encoding windows than the 975 ms used in the long-window analysis. To estimate this encoding window for each channel, we selected the best DNN layer from the long-window analysis and built an encoding model for HGA averaged across 20 ms (-10 to 10 ms relative to the leading edge of the stimulus feature window, which stretches from -975 to 0 ms). If this 20 ms encoding window did not produce a model with R2 > 0.1, then a window of 40 ms was tried. This process was iterated until a sufficiently wide encoding window was found, with candidate windows logarithmically spaced up to 960 ms. Once a sufficient encoding window was identified, separate models were built for each DNN layer using this same window.

Integration windows were then estimated for each channel using the short-window model that maximized R2. Layer activations for the accompanying DNN layer were calculated for spectrograms in which the lagging edge of the spectrogram was truncated to varying degrees, effectively discarding information from earlier timepoints (Fig. 4A, left). For each truncation length, the predicted neural response to all stimuli was estimated and compared to the measured neural responses via correlation. The point at which the predicted response begins to deviate substantially was identified as the integration window; it is at this point that the truncation procedure has begun discarding information that the encoding model has deemed important in modeling the neural response. Quantitatively, this point was identified using the knee point algorithm, which finds the optimal point in which the curve can be split and modeled as two separate line segments. The right panel in Fig. 4A shows two example curves demonstrating short and long integration windows. Integration window results across all patients are shown in Fig. 4B for short-window models with R2 > 0.1. An LME model with the log of the integration window as the response variable revealed an ROI effect, with core containing shorter integration windows than lateral belt (p < 10-5) and parabelt (p = 5.7x10-4; also see Fig. 5, right panel for integration window distributions across ROIs). No significant difference was found between lateral belt and parabelt (p = 0.63) or between left and right hemisphere (p = 0.45).

**Fig. 4.**
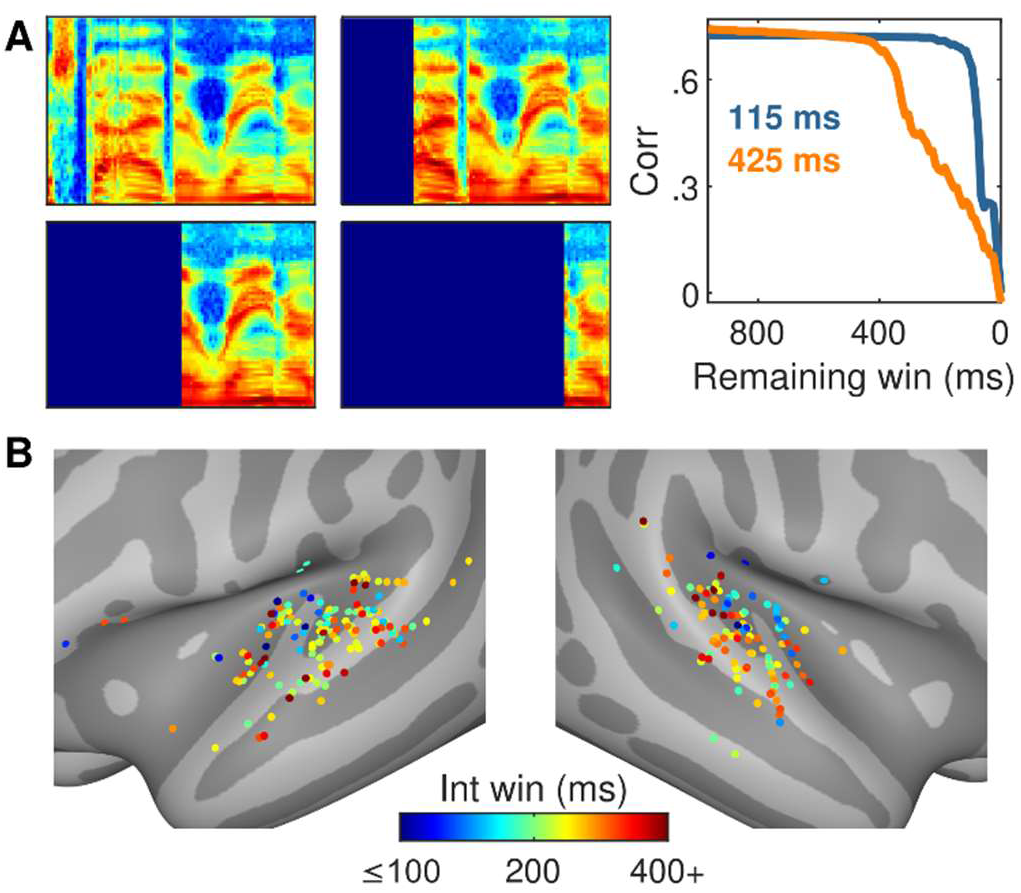
Integration windows. (A) Method for estimating integration windows. Using the best DNN layer’s model for each channel, spectrograms were increasingly truncated and input to the DNN. Predicted HGA was calculated for each truncation window, and correlation was calculated between predicted and observed HGA. The elbow of the correlation curve represents the shortest stimulus window that accurately predicts HGA without appreciable information loss. The right panel shows a core (blue) and lateral belt (orange) channel with integration windows of 115 ms and 425 ms respectively. (B) Integration windows across all patients and channels (with short-window R^2^ > 0.1) are shown, with shorter windows observed in core and longer windows in lateral belt and parabelt regions.

**Fig. 5.**
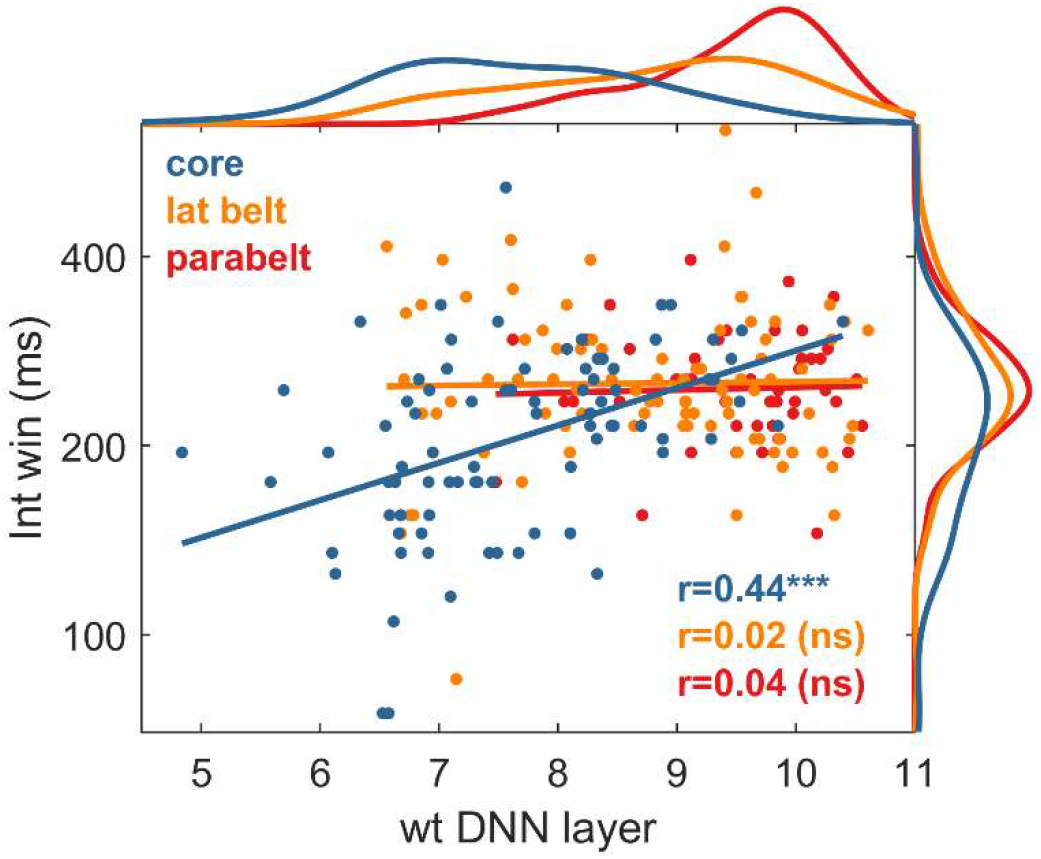
Complexity vs. integration window. The core region showed a strong positive correlation between representational complexity (weighted DNN layer) and integration window length, while lateral belt and parabelt showed no such relationship. Marginal distributions show differences in representational complexity between all three regions (top); integration windows (marginal distribution, right) only differed between core and lateral belt as well as core and parabelt. No differences were observed between lateral belt and parabelt. *** p < 10^-3^, Bonferroni corrected

Lastly, we investigated the relationship between integration windows and representational complexity, shown in Fig. 5. Features of higher complexity often span longer timescales, and neural populations that represent these features likely require longer integration windows to synthesize and code this information. However, as was shown in our previous analysis, both weighted DNN layers and integration windows vary systematically across the auditory cortical hierarchy. To investigate this relationship irrespective of ROI, we built an LME model with the log of the integration window as the response variable and hemisphere, ROI, and weighted DNN layer as fixed effects. An interaction term between ROI and layer was included, and patient was modeled as a random effect. ROI was treated as an ordinal variable with core, lateral belt, and parabelt coded as 1, 2, and 3 respectively. We found significant relationships between weighted DNN layer and integration window, even when controlling for ROI (p < 10-5). ROI also had a significant positive relationship with integration window (p = 5.5x10-5). There was also a significant negative interaction between DNN layer and ROI (p = 1.1x10-4), indicating that the relationship between layer and integration window varied depending on the anatomical region. Specifically, the negative value suggests that the correlation between layer and integration window is strongest in core and decreases with ascending position in the auditory hierarchy. This can be seen in Fig. 5, with a strong positive correlation observed in core and no significant correlations in lateral belt and parabelt regions.

## Discussion

In this work, we demonstrate that neural responses to natural sounds share similarities to representations embedded in a deep neural network (DNN) model trained to categorize sounds. Notably, this DNN accepts acoustic spectrograms as inputs and then outputs a semantic category label, was not exposed to any neural data during its training, and was trained on a separate stimulus set from the one used in this work. Nevertheless, the features that the DNN learned could be used to predict neural responses with high accuracy using an encoding model approach, suggesting an overlap in acoustic representations between the DNN and the brain. Furthermore, sites in increasingly higher-order cortical areas were best predicted by increasingly deeper layers of the DNN, suggesting areas further downstream in the auditory cortical hierarchy represent auditory information with increasing feature complexity. This relationship even existed within specific regions, with smoothly varying gradients of feature complexity observed along relevant anatomical dimensions. Next, we used encoding models to estimate integration windows and found that early auditory areas integrated over shorter windows than higher-order areas. Lastly, we found that channels with increasing complexity used longer integration windows, even when controlling for position in the auditory hierarchy, though this relationship was only observed in early auditory areas.

We showed that auditory cortical responses to natural sounds could be modeled using DNN layer activations and that higher order areas in the auditory hierarchy are best modeled by increasingly deeper layers; both of these findings are aligned with prior studies (15, 24). This relationship between position within the auditory hierarchy and DNN layer depth is consistent with a broadly accepted model in which auditory core, which comprises the posteromedial two thirds of Heschl’s gyrus and is considered to be primary auditory cortex (44, 45), processes low-level acoustic features (2, 46–48). In this model, lateral belt comprises planum temporale along with the anterolateral aspect of Heschl’s gyrus (36, 49) and processes more complex or compositional acoustic features (47, 50–52). Lastly, parabelt is situated in STG and the upper bank of the superior temporal sulcus (STS). While representations in this region are far more heterogeneous, they are best described as high level, abstract features (2, 16, 21, 46, 53).

We also found gradients of complexity within each of core, lateral belt, and parabelt, with these effects lateralizing to the right hemisphere. Within parabelt, complexity increased linearly along a posterior-to-anterior as well as a dorsal-to-ventral dimension. These findings are consistent with other studies that have proposed a flow of information along these axes (45, 54–59). For example, multiple voice processing nodes have been identified in STG and extending into STS (21), with the most posterior node found to process physical speaker characteristics such as vocal tract length (55, 56) and the most anterior one shown to encode individual voice identities (54, 57–59). Notably, these models describe discrete nodes within STG and STS, which would suggest a more step-like transition of complexity along the posterior-anterior axis. However, we found that a linear model was better able to describe the gradient along this axis (as well as the gradient from dorsal to ventral), suggesting a more continuous and smoothly varying degree of complexity. Multiple studies have demonstrated a lateralization of voice processing toward the right hemisphere (21, 57–59), though this finding is sometimes disputed (60). Our measure of complexity was correlated with anatomic position only in the right hemisphere, which may be because conspecific vocalizations represent a particularly salient auditory category, and individual species may have evolved dedicated networks to process it (61). If the shared representations between parabelt sites and DNN layers are largely biased toward voice processing, then this may account for these specific hemispheric differences. Notably, left parabelt/STG is often associated with speech processing (9, 46, 62–64), exhibiting a posterior-to-anterior flow of information (e.g., phonetic features to syllables/words) along the ventral stream (62, 64). While we could still predict left parabelt channels with high accuracy using DNN features, there was no relationship between DNN layer depth and anatomic position within this ROI. This may be because the DNN’s training goals did not require learning progressively compositional speech features, since the speech-related categories it was trained to predict were quite coarse (e.g., speech, babbling, speech synthesizer, etc.).

Notably, we found complexity gradients within the right hemisphere core region. Traditionally, core is believed to represent low-level acoustic features such as relatively simple spectrotemporal representations, typically showing a preference for single frequency bands (48, 65) that are relatively narrow in bandwidth (66, 67). This region is often described as having two subregions, termed hA1 and hR (human homologs to monkey A1 and the rostral area R), which are positioned sequentially along the PM-AL axis. In this model, hA1 and hR exhibit mirrored tonotopic gradients, with hA1 transitioning from high to low frequencies and hR from low to high along the PM-AL dimension (65). Interestingly, our results appear to contradict this model, which would predict a similar representational complexity throughout the core region. One possibility is that this gradient is driven by an increase in integration window length, which would require deeper layers to model within the DNN, given the narrow 3x3 extent of the DNN’s convolutional filters. This possibility follows from the observation in primates that the core subregion R integrates over longer windows than A1 (68). However, this model would predict a stepwise or sigmoidal relationship across the hA1-hR boundary, which is inconsistent with our findings of a linear relationship.

Lastly, lateral belt areas have been shown to respond to more complex sounds such as band-passed noise (47, 51). This area is also parcellated into multiple subregions, including hML and hAL (human homologs to middle lateral and anterolateral belt regions), which are positioned sequentially along the PM-AL axis. Studies have found that hML contains a tonotopic gradient, suggesting relatively lower-level acoustic processing, while hAL overlaps with voice-sensitive regions and exhibits a strong preference for low frequencies (a prominent feature of human vocalizations) (36, 67, 69). This latter finding suggests that hAL may engage in higher-order processing to support complex or abstract representations of voice (67). The complexity gradient we observed along the PM-AL axis of lateral belt is consistent with these findings, including the fact that this gradient was better modeled by a sigmoidal (i.e., step-like) relationship compared to a linear one. It should be noted that for all analyses comparing linear and sigmoidal relationships for complexity gradients within ROIs, it is possible that these findings may be impacted by spatial blurring due to factors such as individual differences in anatomy and volume conduction; nevertheless, we believe they represent intriguing findings that warrant further investigation in future studies.

Consistent with other lines of evidence, our short-window encoding models found that early auditory cortex (core) integrates over shorter windows compared to downstream regions in the auditory hierarchy (39, 70, 71). In contrast with previous work (39), we did not observe differences between non-primary areas of the hierarchy (i.e., lateral belt and parabelt). This could be attributed to a lack of sensitivity in the estimation of integration windows: encoding models may overestimate the temporal extent of the encoded features, given the statistical regularities and local correlations inherent in natural sounds. Nevertheless, our approach was able to identify relative differences in integration windows between primary and non-primary regions.

Lastly, we found that channels that represented more complex features had longer integration windows. Importantly, this effect persisted even when accounting for gross anatomical location (ROI position within the auditory hierarchy); this control is necessary, since our results and other studies have shown that integration windows increase across the auditory hierarchy (39, 70, 71). Further exploration revealed that this relationship was observable in core but not in lateral belt or parabelt. It is unclear whether this apparent regional specificity is a genuine property or is an artefact of the sensitivity issues mentioned previously (i.e., temporally local correlations within natural sounds could potentially lead to an overestimation of integration window length). Another possibility is that there is relatively less variability in the estimated integration windows in non-primary areas relative to core, as well as less variability in weighted DNN layers for parabelt compared to core and lateral belt; these factors may provide less opportunity for covariation between integration windows and representational complexity in higher-order areas. Given this ambiguity, we will avoid making inferences on the absence of a relationship in non-primary areas. However, the significant correlation observed within core channels does support the general principle that more complex neural representations require longer time windows to integrate and combine simpler features into higher-order compositional representations, likely due (at least in part) to the longer temporal extent that these representations span. Given the caveat that our results only show this effect in primary auditory cortex, future studies would be required to identify whether this principle applies to higher-order as well as other primary sensory cortical regions.

In considering the findings presented here, several caveats should be taken into account. First, as with nearly all studies involving intracranial recordings, our electrodes are implanted based on clinical necessity, which may cause a selection bias for recordings sites. While the patients involved are not neurotypical, their perceptual performance often is, and the spatial and temporal resolution of these direct cortical recordings provide substantial value both in testing existing theories and offering new findings that can then be tested using other experimental modalities and approaches. Furthermore, the large number of patients included in this study (N = 20) and the heterogeneity of their pathologies (see Table 1) provide some assurance that any individual deviations would have minimal impact on the group results, assuming that those deviations are distributed randomly. Another caveat related to the nature of the recordings is the sparsity of sEEG recordings, both within individual regions and across hemispheres (only a subset of patients has bilateral implantations). This limitation would have the effect of reducing statistical power, and thus positive statistical results can be considered with this in mind. Lastly, throughout the paper we have interpreted the DNN layer depth as a measure of representational complexity. While layer depth may seem like a rough proxy for this concept, it is a direct measure of the number of nonlinear transformations the input has undergone (26). Thus, we believe it is a straightforward and readily interpretable index of complexity, especially considering the way it has neatly mapped on to existing models of auditory cortex.

## Abbreviations

DNN: deep neural network
HGA: high-gamma activity
LME: linear mixed effects
ROI: region of interest
sEEG: stereo-electroencephalography
STM: spectrotemporal modulation
STG: superior temporal gyrus
STS: superior temporal sulcus
STP: supratemporal plane

